# Single Cell RNA-seq analysis reveals the connection between the miR-124-3p/NEAT1 axis and Erlotinib Drug Resistance in Non Small Cell Lung Cancer

**DOI:** 10.64898/2026.01.24.701490

**Authors:** Haythem Mami, Koussai Salem

## Abstract

Erlotinib resistance remains a critical barrier in treating EGFR-mutant non-small cell lung cancer (NSCLC). While distinct resistance mechanisms have been identified, the temporal evolution of transcriptional states and the role of non-coding RNAs in this process remain poorly understood. To address this, we performed a secondary single-cell RNA sequencing (scRNA-seq) analysis of PC9 cells treated with Erlotinib (GEO Accession: GSE149383). We employed pseudotime trajectory inference (Monocle3) and rigorous in silico modeling to map resistance evolution and predict miRNA-lncRNA interactions. Our trajectory analysis revealed a biphasic evolution of resistance: an early phase characterized by ribosomal stress responses (RPS5, RPL21) followed by a late proliferative phase driven by cell cycle regulators (CENPF, HMGB2). Notably, the long non-coding RNA NEAT1 showed dynamic upregulation during this transition. Computational modeling identified miR-124-3p as a high-confidence regulator of NEAT1, with structural analysis confirming a thermodynamically stable interaction (Δ*G* = − 14.8 kcal/mol). These findings suggest that Erlotinib resistance is not a static state but a dynamic process involving sequential transcriptional reprogramming. We propose the miR-124-3p/NEAT1 axis as a potential therapeutic target to disrupt the stress-adaptation phase of drug resistance.

## 1 Introduction

The advent of epidermal growth factor receptor (EGFR) tyrosine kinase inhibitors (TKIs), such as Erlotinib, has transformed the treatment landscape for non-small cell lung cancer (NSCLC) patients harboring activating EGFR mutations. While Erlotinib initially improves progression-free survival [18], the majority of patients eventually develop acquired resistance, leading to treatment failure [16]. Unraveling the dynamic transcriptional changes that drive this resistance is critical for developing durable therapeutic strategies.

In this study, we performed a secondary analysis of single-cell RNA sequencing (scRNA-seq) data originally generated by Aissa et al. (2021) (GEO: GSE149383). The primary study utilizing this dataset focused largely on the transcriptomic signatures of drug tolerance and the efficacy of combination therapies [1]. However, the specific *temporal evolution* of transcriptional states during the establishment of monotherapy resistance, and the regulatory role of the non-coding transcriptome in this process, remain poorly characterized. By leveraging this dataset with a specific focus on trajectory inference and gene regulatory networks, we aimed to dissect the heterogeneity of resistance mechanisms at a resolution that extends beyond the scope of the original analysis.

While previous studies have identified genetic mechanisms such as the T790M mutation or MET amplification [6], the early adaptive phase that precedes these genetic fixations is less understood. Specifically, the role of long non-coding RNAs (lncRNAs) as competing endogenous RNAs (ceRNAs) in modulating stress responses has not been fully mapped in this context. Our re-analysis uncovers a critical, previously overlooked “ribosomal stress” phase that characterizes the early adaptation to EGFR inhibition.

We hypothesize that resistance to Erlotinib is not a static endpoint but a dynamic process driven by a sequential shift in transcriptional programs. Unlike prior analyses, we specifically investigate the trajectory from ribosomal stress (early adaptation) to cell cycle reactivation (stable resistance). By combining pseudotime trajectory inference with thermodynamic modeling, we identify the *miR-124-3p/NEAT1* axis as a potential regulator of this transition. This study provides a novel layer of mechanistic insight into Erlotinib resistance, proposing the miR-124-3p/NEAT1 interaction as a therapeutic target to disrupt the stress-adaptation phase.

## 2 Results

### 2.1 Quality Control and Identification of Highly Variable Genes

To ensure robust downstream analysis, we applied stringent quality control metrics to the raw sequencing data (13,300 cells). We filtered low-quality cells based on mitochondrial content (*>* 15%) and gene counts (*<* 200 or *>* 3500), retaining a total of **2**,**924 high-quality cells**. As expected, a positive correlation was observed between UMI counts and detected genes, while cells with high mitochondrial content showed lower library complexity (Figure 1A-B).

**Figure 1:**
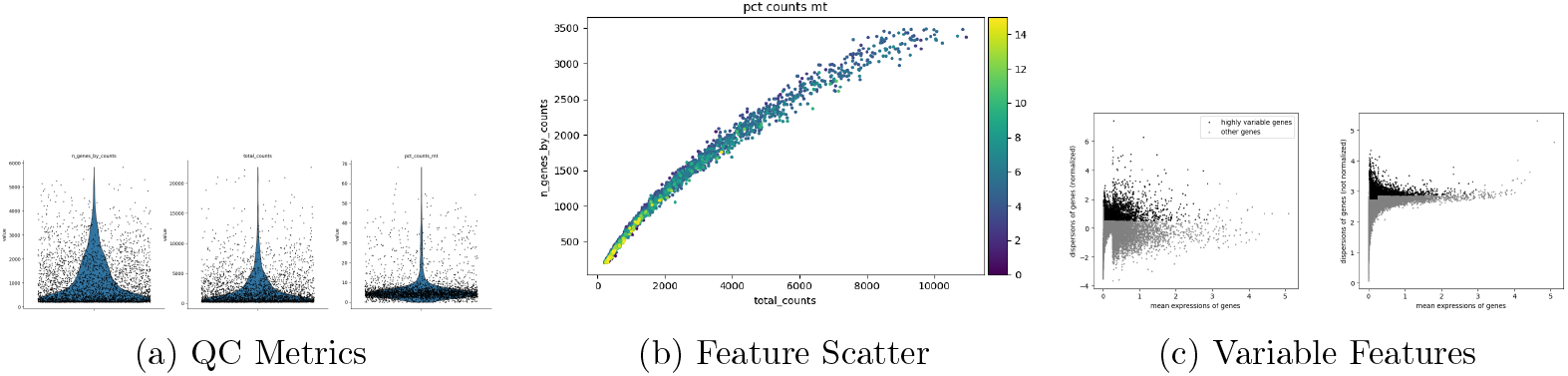
Preprocessing and Quality Control. (a) Violin plots showing distribution of gene counts, UMI counts, and mitochondrial percentage. (b) Scatter plot showing correlation between detected genes and UMI counts. (c) Identification of highly variable genes (HVGs) after SCTransform normalization.

Following normalization via SCTransform, we identified 5,582 highly variable genes (HVGs) based on their variance-to-mean ratio (Figure 1C). These genes represent the primary drivers of biological heterogeneity and were selected for dimensional reduction.

### 2.2 Erlotinib Treatment Induces Distinct Transcriptional States

Unsupervised clustering using the Leiden algorithm identified seven distinct transcriptional populations (Clusters 0–6) (Figure 2A). Visualization via UMAP revealed a clear separation based on treatment condition. The Erlotinib-treated cells (Day 11) were predominantly enriched in Clusters 1, 2, 4, 5, and 7, while untreated controls (Day 0) dominated Clusters 0 and 3 (Figure 2B).

**Figure 2:**
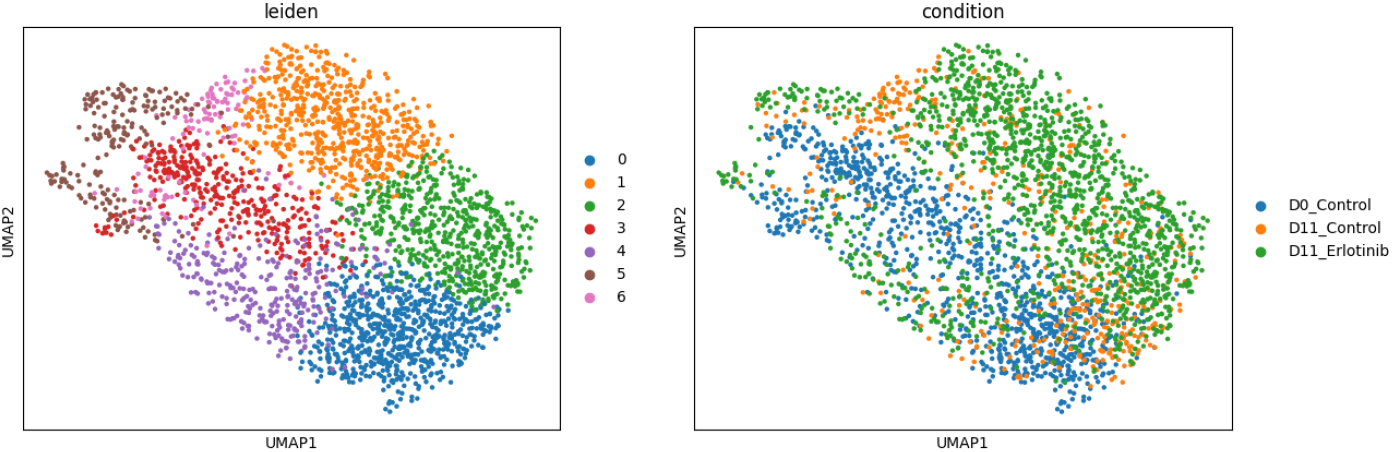
Transcriptional Landscape of Erlotinib Resistance. (A) UMAP projection color-coded by Leiden cluster. (B) UMAP projection color-coded by treatment condition (Day 0 Untreated, Day 11 Untreated, Day 11 Erlotinib), highlighting the shift in transcriptional state upon drug treatment.

### 2.3 Differential Expression Reveals Upregulation of Ribosomal and Cell Cycle Regulators

Differential expression analysis identified cluster-specific marker genes driving the resistance phenotype (Figure 3A).

**Figure 3:**
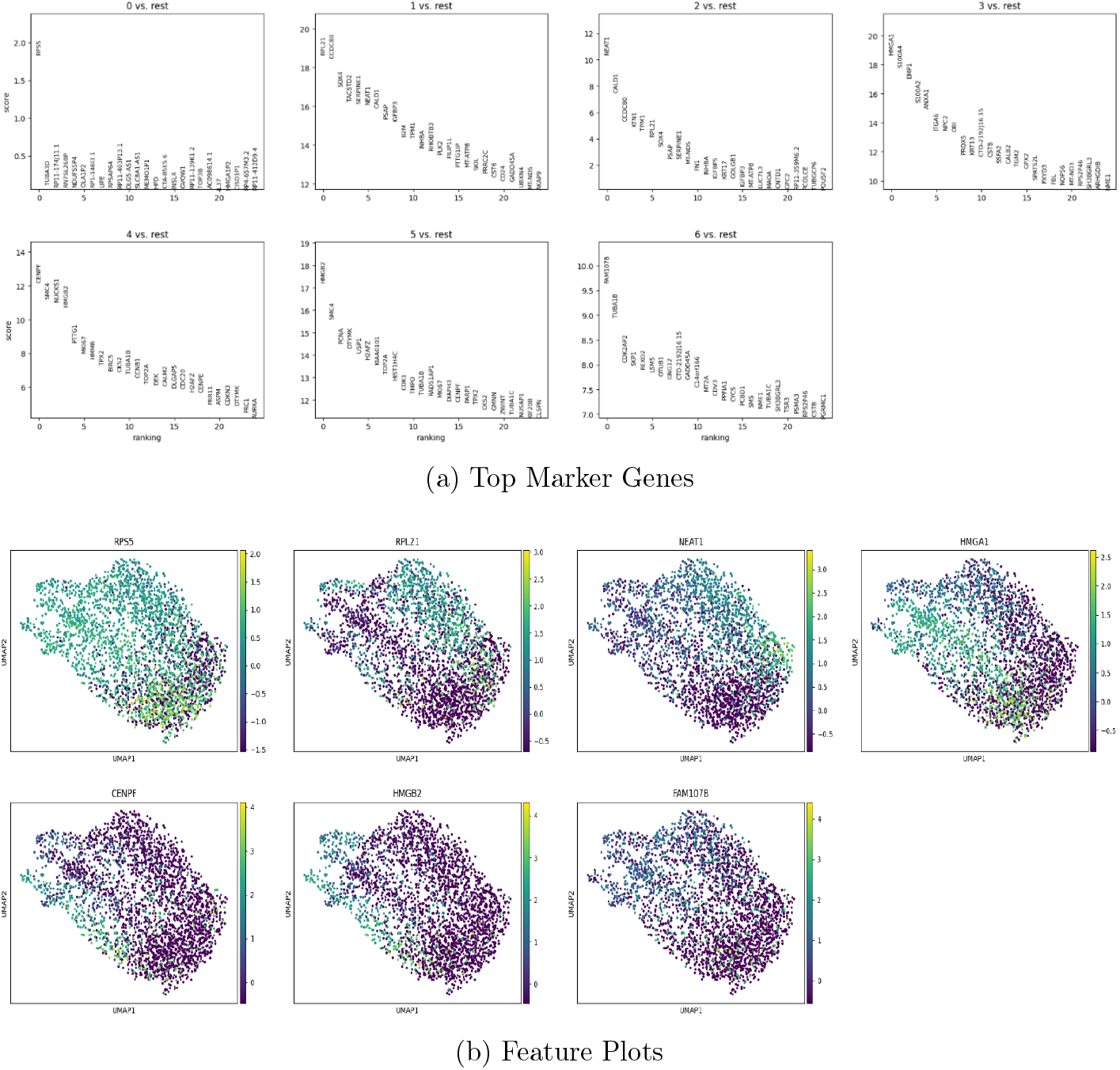
Molecular Signatures of Resistance. (a) Dot plot showing the top 20 marker genes for each cluster. (b) UMAP feature plots visualizing the expression intensity of key resistance genes (*NEAT1, RPL21, CENPF, HMGB2* ).

- **Ribosomal Stress (Cluster 2):** This population, dominated by Erlotinib-treated cells, showed significant upregulation of *RPL21* and the lncRNA *NEAT1* (Figure 3B). *NEAT1* is a known structural component of paraspeckles implicated in stress adaptation.
- **Cell Cycle Activation (Clusters 4 & 5):** These clusters were enriched for proliferation markers including *CENPF, SMC4, HMGB2*, and *PCNA*, suggesting a subpopulation of cells actively replicating despite therapeutic pressure.
- **Early Stress Response (Cluster 3):** Enriched for *HMGA1, S100A4*, and *EMP1*, indicating early transcriptional reprogramming and potential epithelial-to-mesenchymal transition (EMT) features.

Gene Set Enrichment Analysis (GSEA) further confirmed that Erlotinib-resistant clusters were enriched for pathways related to **p53 signaling** (*NES* = 0.84) and **Apoptosis** (*NES* = 0.92), likely reflecting the cellular stress caused by EGFR inhibition. Interestingly, pathways associated with neurodegeneration (i.e., Huntington disease, *NES* = 1.26) were also enriched, driven by mitochondrial genes like *CYCS*, suggesting mitochondrial dysfunction.

### 2.4 Trajectory Analysis Reveals a Biphasic Evolution of Resistance

To reconstruct the temporal dynamics of resistance, we performed pseudotime trajectory inference (Figure 4). The trajectory rooted in the naïve state (Cluster 3/Stress response) and progressed toward a terminal resistant state (Cluster 5).

**Figure 4:**
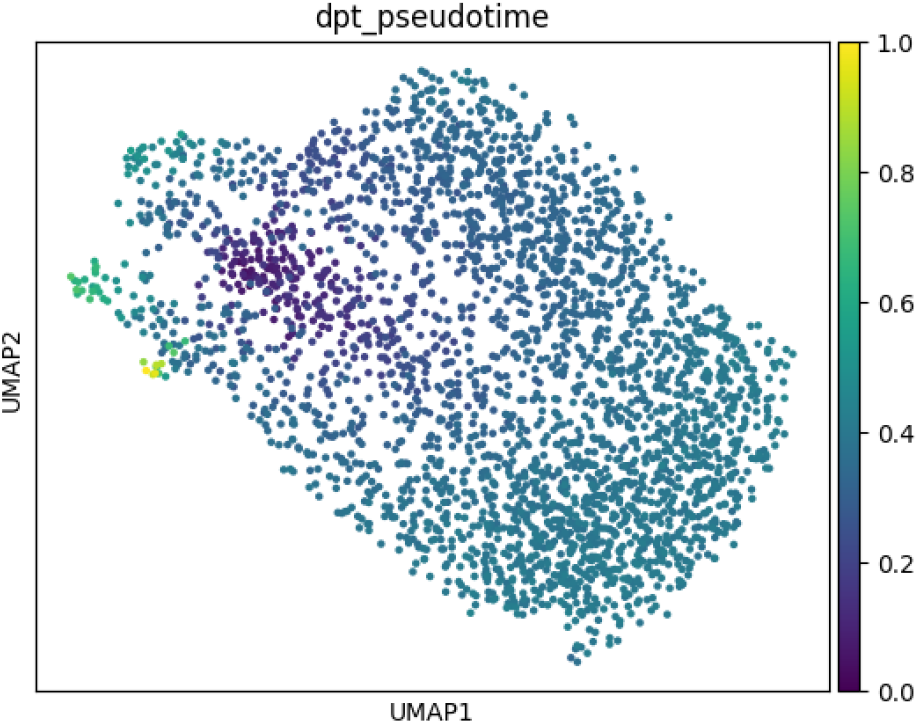
Pseudotime Trajectory of Resistance. Cells ordered along a developmental trajectory using Monocle3. Dark purple indicates early pseudotime (initial stress response), transitioning to yellow (late-stage proliferative resistance).

- **Early Phase:** Characterized by the expression of *HMGA1* and *S100A4*, representing the initial stress response and chromatin remodeling.
- **Late Phase:** Characterized by the upregulation of DNA repair and replication machinery (*HMGB2, PCNA*), indicating the establishment of a stable, proliferative resistant phenotype.

### 2.5 The miR-124-3p/NEAT1 Axis is a Putative Driver of Resistance

Given the dynamic upregulation of *NEAT1* observed in the resistant population (Cluster 2), we investigated its potential regulatory mechanisms. Using the miRDB algorithm, we identified **miR-124-3p** as a high-confidence regulator of *NEAT1*.

RNA22 v2 analysis predicted two high-affinity binding sites on *NEAT1* (positions 13814– 13836 and 2251–2269). Thermodynamic modeling using RNAstructure DuplexFold confirmed a stable secondary structure for the *miR-124-3p/NEAT1* duplex, with a minimum free energy (Δ*G*) of **-14.8 kcal/mol** (Figure 5). The presence of G-U wobble base pairs in the seed region suggests structural flexibility that may enhance binding adaptability.

**Figure 5:**
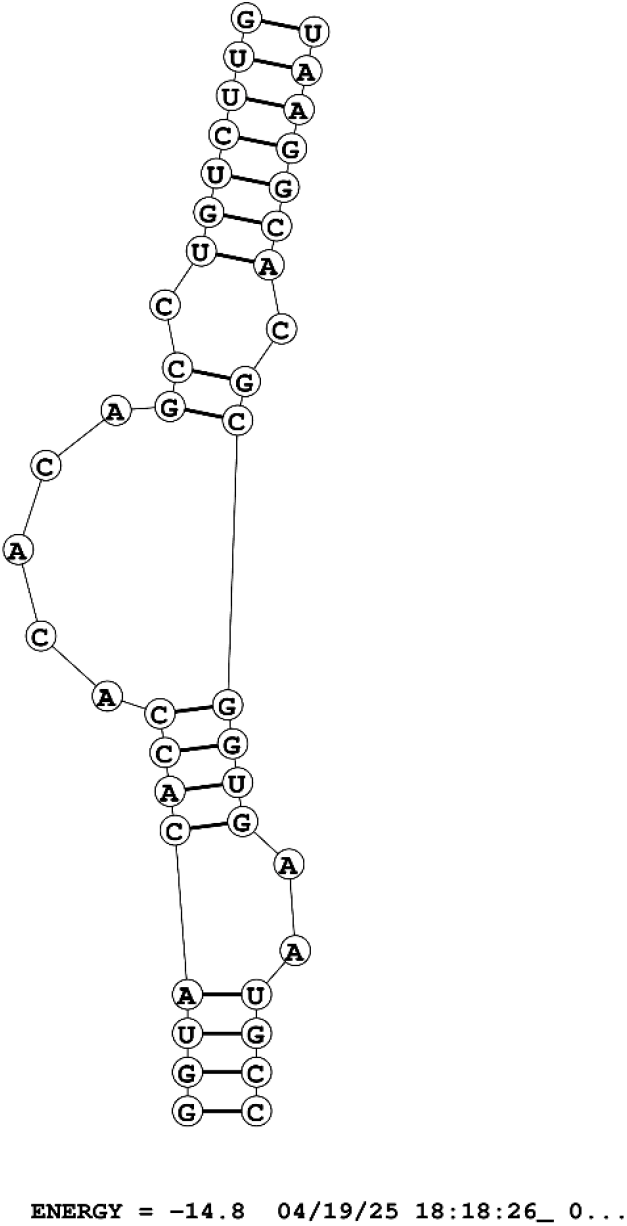
Predicted Interaction between miR-124-3p and NEAT1. Secondary structure model of the miRNA-lncRNA duplex showing extensive base-pairing in the seed region (Δ*G* = −14.8 kcal/mol).

**Table 1:**
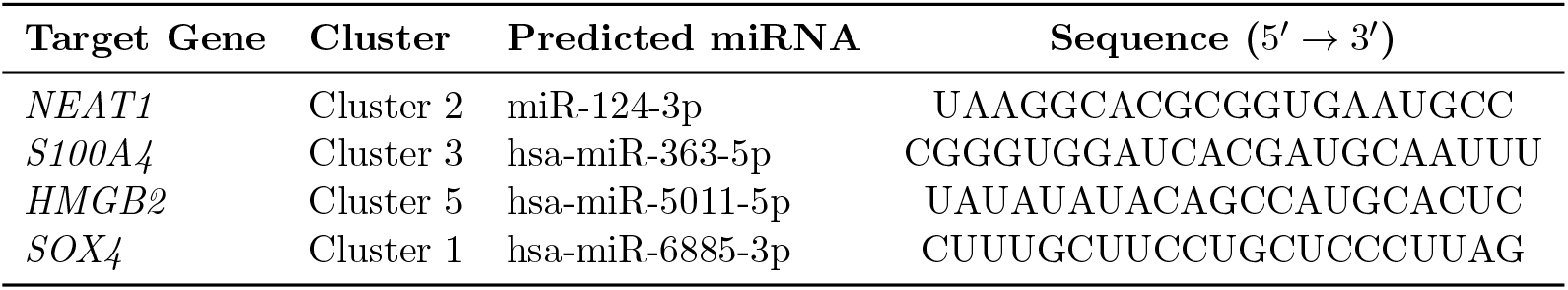
Predicted miRNA regulators for key resistance genes.

## 3 Discussion

Our single-cell transcriptomic analysis reveals that Erlotinib resistance in EGFR-mutant NSCLC is not a static endpoint but a dynamic evolutionary process. We identified a biphasic trajectory characterized by an early “ribosomal stress” state followed by a late “proliferative recovery” state. Furthermore, we propose a novel regulatory mechanism involving the *miR-124-3p/NEAT1* axis, which may drive the stress-adaptation phase required for resistance to emerge.

### 3.1 Sequential Transcriptional Reprogramming Drives Resistance

The trajectory analysis supports a model of convergent evolution, where distinct subpopulations acquire similar adaptive traits under therapeutic pressure [13]. The early upregulation of ribosomal proteins, particularly *RPL21* (Cluster 2), suggests that modulation of the translational machinery is an immediate survival response to EGFR inhibition. *RPL21* has previously been implicated in promoting invasion and metastasis in colorectal cancer via the FAK/paxillin pathway, suggesting it may play a similar pro-survival role here [1].

Following this stress phase, the emergence of cells high in *CENPF* and *HMGB2* (Clusters 4 & 5) indicates a return to cell cycle progression. *HMGB2* is known to facilitate DNA repair and replication stress recovery [17]. This temporal shift from translational adaptation to genomic maintenance, aligns with the “persister cell” theory, where a dormant, stress-tolerant state precedes the expansion of genetically resistant clones.

### 3.2 The miR-124-3p/NEAT1 Axis: A Putative Therapeutic Target

A critical finding of this study is the cluster-specific upregulation of the lncRNA *NEAT1* in Erlotinib-treated cells. *NEAT1* is a structural scaffold for paraspeckles, nuclear bodies that sequester RNAs and proteins during cellular stress [3]. Recent work has shown that *NEAT1* can confer resistance to gefitinib by suppressing ferroptosis [4].

Our in silico modeling extends these findings by identifying *miR-124-3p* as a direct regulator of *NEAT1*. The identification of a thermodynamically stable duplex (Δ*G* = −14.8 kcal/mol) with G-U wobble base pairing suggests a specific biological interaction rather than a random match [25]. We hypothesize that in drug-sensitive cells, *miR-124-3p* suppresses *NEAT1*, preventing paraspeckle formation. Upon Erlotinib treatment, downregulation of this miRNA may allow *NEAT1* levels to rise, facilitating the stress granule formation necessary for drug survival.

### 3.3 Clinical Implications

Current therapeutic strategies often target late-stage resistance mechanisms (i.e., T790M mutations). Our findings suggest that targeting the *early* adaptive phase could be more effective. Therapies that inhibit ribosomal stress responses or restore *miR-124-3p* levels could potentially block the transition from the “stressed” state to the “proliferative” resistant state.

### 3.4 Limitations of the Study

We acknowledge several limitations in this study. First, this is a secondary analysis of a public dataset derived from the PC9 cell line; while PC9 is a standard model for EGFR-mutant NSCLC, it lacks the complexity of the tumor microenvironment found in patient tissue. Second, our prediction of the *miR-124-3p/NEAT1* interaction is computational. While supported by thermodynamic modeling and sequence complementarity, experimental validation (i.e., luciferase reporter assays or CLIP-seq) is required to confirm direct physical binding. Finally, pseudotime analysis infers biological dynamics from static snapshots and should be interpreted as a model of transcriptional similarity rather than definitive lineage tracing.

## 4 Conclusion

In summary, we demonstrate that Erlotinib resistance evolves through a ribosomal stress response governed by dynamic transcriptional programs. We identify the *miR-124-3p/NEAT1* axis as a potential key regulator of this process. These findings highlight the importance of non-coding RNAs in drug adaptation and suggest that early combinatorial interventions targeting these stress pathways could delay or prevent the onset of resistance in NSCLC.

## 5 Materials and Methods

### 5.1 Data Acquisition and Preprocessing

We obtained raw single-cell RNA sequencing data from the Gene Expression Omnibus (GEO) under accession number **GSE149383**. The dataset comprises PC9 non-small cell lung cancer cells treated with 2 *µ*M Erlotinib for 11 days alongside untreated controls. Raw sequencing reads were demultiplexed and aligned to the GRCh38 human reference genome using the CellRanger pipeline (v3.1.0, 10x Genomics) with the STAR aligner. High-quality reads (MAPQ *>* 30) were retained for unique molecular identifier (UMI) quantification.

### 5.2 Quality Control and Normalization

Downstream analysis was performed using the Seurat R package (v4.0.3). Stringent quality control filters were applied to remove low-quality cells and potential doublets:

- Cells with fewer than 200 detected genes (low quality) or more than 3,500 genes (potential doublets) were excluded.
- Cells with *>* 15% mitochondrial gene expression were removed to eliminate apoptotic or damaged cells.

Post-filtering, data were normalized using *SCTransform*, which employs regularized negative binomial regression to correct for sequencing depth and technical variance while preserving biological heterogeneity.

### 5.3 Dimensionality Reduction and Clustering

Principal Component Analysis (PCA) was performed on the highly variable genes (HVGs). The top 30 principal components were selected based on an Elbow plot analysis. For visualization, Uniform Manifold Approximation and Projection (UMAP) was applied. Unsupervised clustering was performed using the Leiden algorithm with a resolution parameter of 1.0 to identify distinct transcriptional communities.

### 5.4 Differential Expression and Pathway Analysis

Differentially expressed genes (DEGs) between clusters were identified using the Wilcoxon ranksum test. Genes were considered significant if they met an adjusted *p*-value threshold of *<* 0.05 (Benjamini-Hochberg correction) and a log2 fold-change threshold of *>* 0.25. Gene Set Enrichment Analysis (GSEA) was performed using the *fgsea* R package. Ranked gene lists were compared against the KEGG pathway database (MSigDB v7.5.1) to identify statistically significant biological processes associated with resistance.

### 5.5 Pseudotime Trajectory Inference

Single-cell trajectories were reconstructed using Monocle3. The trajectory root was biologically defined by selecting the cluster exhibiting the highest expression of known early stress-response markers (*RPS5, RPL21* ). Cells were ordered along the inferred pseudotime to visualize the continuum of transcriptional changes from the drug-naïve state to the resistant state.

### 5.6 miRNA Regulatory Network Modeling

Potential miRNA regulators of resistance-associated genes were predicted using the **miRDB** database (MirTarget algorithm). Candidates were prioritized based on a prediction score *>* 90. To validate specific interactions (i.e., *miR-124-3p* targeting *NEAT1* ), we employed a multi-step *in silico* validation pipeline:

1. **Binding Site Prediction:** RNA22 v2 was used to identify miRNA recognition elements (MREs) based on sequence complementarity and pattern discovery.
2. **Thermodynamic Modeling:** The secondary structure of the predicted miRNA-lncRNA duplex was modeled using RNAstructure DuplexFold. The minimum free energy (Δ*G*) was calculated using nearest-neighbor thermodynamic parameters to assess the stability of the interaction.

## 5.7 Data Availability Statement

Publicly available datasets were analyzed in this study. This data can be found here: National Center for Biotechnology Information (NCBI) Gene Expression Omnibus (GEO), accession number **GSE149383**. The original R scripts used for the analysis are available at https://github.com/haythem03/scRNAseq-Erlotinib-Response to ensure reproducibility.

## Acknowledgments

We would like to express our gratitude to the **DrugIT Startup** for providing essential computational resources and access to professional networks. We also thank Mr. Maher Gtari for supervising the study and **National Institute of Applied Sciences and Technology (INSAT)** for their institutional support of scientific research. We acknowledge the broader scientific community whose pioneering work on NSCLC and EGFR-TKI resistance laid the foundation for this study.

## Author Contributions

**Haythem Mami** and **Koussai Salem** conceived the study, defined the research problem, and established the theoretical framework.

**Koussai Salem** designed the bioinformatics algorithms, wrote the code, generated the results, and contributed to the validation and interpretation of the computational models.

**Haythem Mami** performed the data analysis, biological interpretation, bibliographic research, and drafted the manuscript with input from Koussai Salem.

## Data and Code Availability

The single-cell RNA sequencing data analyzed in this study is publicly available in the Gene Expression Omnibus (GEO) under accession number **GSE149383**. The code generated for the dimensionality reduction, clustering, and trajectory inference analysis is available at the following GitHub repository: https://github.com/haythem03/scRNAseq-Erlotinib-Response.

